# MealTime-MS: A Machine Learning-Guided Real-Time Mass Spectrometry Analysis for Protein Identification and Efficient Dynamic Exclusion

**DOI:** 10.1101/2020.05.22.110726

**Authors:** Alexander R. Pelletier, Yun-En Chung, Zhibin Ning, Nora Wong, Daniel Figeys, Mathieu Lavallée-Adam

## Abstract

Mass spectrometry-based proteomics technologies are the prime methods for the high-throughput identification of proteins in complex biological samples. Nevertheless, there are still technical limitations that hinder the ability of mass spectrometry to identify low abundance proteins in complex samples. Characterizing such proteins is essential to provide a comprehensive understanding of the biological processes taking place in cells and tissues. Still today, most mass spectrometry-based proteomics approaches use a data-dependent acquisition strategy, which favors the collection of mass spectra from proteins of higher abundance. Since the computational identification of proteins from proteomics data is typically performed after mass spectrometry analysis, large numbers of mass spectra are typically redundantly acquired from the same abundant proteins, and little to no mass spectra are acquired for proteins of lower abundance. We therefore propose a novel supervised learning algorithm that identifies proteins in real-time as mass spectrometry data are acquired and prevents further data collection from confidently identified proteins to ultimately free mass spectrometry resources to improve the identification sensitivity of low abundance proteins. We use real-time simulations of a previously performed mass spectrometry analysis of a HEK293 cell lysate to show that our approach can identify 92.1% of the proteins detected in the experiment using 66.2% of the MS2 spectra. We also demonstrate that our approach outperforms a previously proposed method, is sufficiently fast for real-time mass spectrometry analysis, and is flexible. Finally, MealTime-MS’ efficient usage of mass spectrometry resources will provide a more comprehensive characterization of proteomes in complex samples.

## INTRODUCTION

Comprehensive identification of proteins in complex samples is best achieved by mass spectrometry (MS)-based proteomics techniques^1–4^. Among these proteomics approaches, we count bottom-up proteomics, which performs a tryptic digestion of the proteins in a sample into peptides, followed by a chromatographic separation of those peptides, and their ionization, such that they can be analyzed by a mass spectrometer, where the mass-over-charge (m/z) values of the ionized peptides are measured and quantified. Such an approach uses tandem mass spectrometry (MS/MS) to measure the m/z value of precursor ions (MS1 spectrum) and collects mass spectra (MS2 spectra) of fragmented peptides, which are then used for peptide sequence determination and protein identification. Since not all ions produced in an experiment can be fragmented to generate MS2 spectra due to time constraints, different mass spectra acquisition strategies have been proposed to maximize the number of protein identifications^5–9^. While technological improvements have accelerated mass spectrometry analysis and improved peptide separation to allow the acquisition of more MS2 spectra from the proteins in a sample^10–15^, the field is still far from a comprehensive MS2 spectrum acquisition covering all peptide species present in an experiment.

### Top N data-dependent acquisition

One of the most popular approaches for MS2 spectra acquisition is called Top N Data-Dependent Acquisition (DDA). This method collects MS2 spectra for the N most intense ions in an MS1 spectrum. These ions typically correspond to peptides belonging to the most abundant proteins in a sample. They are also more likely to generate MS2 spectra that have a higher signal-to-noise ratio and that are often easier to identify than those generated from ions with lower intensities. Acquiring mass spectrometry data with this strategy favors the analysis of peptides of high abundance, while obtaining little to no MS2 spectra for peptides of lower abundance, thereby limiting the protein identification sensitivity for the experiment^16,17^. In fact, high abundance proteins are often analyzed in excess of what is needed for a confident protein identification with this approach.

### Dynamic Exclusion to Improve Protein Identification Sensitivity

As a result of liquid chromatography (LC) separation, a given peptide species is only present for a given elution time window during an MS experiment. However, peptides originating from high abundance proteins can elute over long periods of time. These long elution time periods, will cause DDA approaches to redundantly analyze the same highly abundant peptides, while gathering little to no data related to less abundant ones. To resolve this issue, a common strategy is to employ a dynamic exclusion approach. Dynamic exclusion prevents the fragmentation of ions with a given m/z value for a short period of time after an MS2 spectrum was acquired for that same m/z value^18^. The mass spectrometer therefore dynamically maintains a list, over time, of m/z values for which ions are excluded from fragmentation, thus giving the opportunity for ions of peptides of lower intensity to be analyzed. Nevertheless, since abundant proteins typically generate numerous peptides with different charge states and that abundant peptides can elute over periods of time much longer than their exclusion periods (typically 30 to 90 seconds^19^), mass spectrometry DDA approaches still acquire more MS2 spectra than necessary for their confident identification despite this exclusion strategy.

### Alternative MS/MS exclusion strategy

In addition to traditional dynamic exclusion, other exclusion strategies have been proposed to increase protein identification sensitivity of MS/MS. Among these, we count the Bespoke exclusion list approach, which excludes peptides from fragmentation based on a pre-defined list of high abundance and expected analytes that are not informative in a typical proteomics experiment^19^. Such an exclusion list often includes protein contaminants, such as keratins, proteolytic enzymes used in sample preparation, and small molecules, such as organic solvents and detergents used in lab equipment cleaning. While this approach improves MS data acquisition by reducing the analytical time devoted to contaminants that represent experimental artefacts, high abundance proteins that are biologically or experimentally relevant remain likely to prevent the acquisition of MS2 spectra from peptides of low abundance.

Kreimer et al. also addressed low abundance protein identification issues with their SmartMS algorithm^20^. Their strategy uses an exclusion list to prevent the selection of m/z values likely corresponding to peptides that have been confidently identified in a number of previously performed technical replicate experiments with the goal of increasing the number of confidently identified proteins. After each technical replicate MS analysis of aliquoted samples, confidently identified peptides are added to the exclusion list of the following replicate’s MS analysis. Although Kreimer et al. demonstrated that their approach was successful in identifying more proteins when compared to conventional Top N DDA^20^, their algorithm relies on repeated analysis of a sample and thus requires samples where quantity is not a problem. It also consumes lots of MS resources to iteratively exclude highly abundant peptides after repeated experiments.

### Pseudo real-time and real-time mass spectrometry analyses

An alternative to excluding ions from fragmentation after the completion of an MS experiment is to identify peptides and proteins on-the-fly during MS analysis and to update the exclusion list in real-time based on those identifications. McQueen et al. proposed an approach working towards that goal^21^. Their method periodically updates an exclusion list based on proteins it has identified so far in the experiment. This update is made at a small number of chromatographic time points using spiked-in peptides with known liquid chromatography retention times. Although this approach effectively reduces the redundant analysis of peptides from proteins identified with high confidence, the periodic update of the exclusion list does not fully take advantage of the information available in real-time, as peptides are only added to the exclusion list at the following pre-determined time point, which can occur several minutes after a protein is confidently identified. Additionally, their approach uses fixed heuristic thresholds based on spectral counts and protein identification confidence scores for assessing the reliability of a confident protein identification. The optimal values for these thresholds can be difficult to determine and unfortunately any peptides with scores below these thresholds do not contribute to determining a protein’s identification confidence.

Recently, a number of approaches using real-time analysis of MS data have been proposed. For instance, the Real Time Search-MS3 algorithm identifies peptides in real-time and assesses the confidence of their identification in order to decide whether it is worth acquiring an MS3 spectra for quantification purposes^22^. In other words, the method only tries to quantify peptides with an MS3 spectra if their parent MS2 spectra is identifiable. Another method, named Advanced Peak Determination, uses an iterative approach to detect isotopic clusters that have similar m/z values^23^. The approach provides a sophisticated peak determination strategy that increases peptide and, thereby, protein identification sensitivity. Finally, the MaxQuant.Live algorithm increases the number of peptides that can be quantified using targeted MS^24^. The algorithm recognizes in real-time peptides from MS1 spectra based on a targeted list of peptides a user aims to quantify. It does so by using the peptides’ m/z value, retention time, and intensity. These real-time MS methods show the wide range of applications of processing mass spectrometry data in realtime to characterize peptides and control the instrument’s behavior. However, to our knowledge, no approaches are dealing with the problem of identifying proteins in real-time during mass spectrometry experiments with the goal of increasing the identification sensitivity of low abundance proteins. Furthermore, machine learning strategies have been severely underexploited to analyze MS data in real-time.

Herein, we present a novel machine-learning-guided realtime mass spectrometry approach that identifies proteins on-the-fly during DDA and that uses these protein identifications to build in real-time a dynamic exclusion list. Using real-time simulations from a previously acquired MS dataset, we show that our software package, named MealTime-MS, can confidently identify the vast majority of proteins of a traditional DDA experiment (92.1%), while using only a fraction of the spectra (66.2%). We also show that our supervised learning algorithm outperforms an approach inspired by McQueen et al.’s work using heuristic determination of protein identification^21^. In contrast with their method, MealTime-MS’ flexibility can be controlled with a simple parameter adjustment. Finally, we demonstrate that MealTime-MS is sufficiently fast to efficiently analyze MS data in real-time.

## METHODS

### Method overview

We propose a machine learning approach, MealTime-MS, for the real-time identification of proteins during MS experiments. A logistic regression classifier uses the results of a sequence database search to assess the confidence of protein identifications. MealTime-MS then prevents data acquisition of mass spectra from peptides that belong to already confidently identified proteins by maintaining a dynamic exclusion list in real-time. Figure 1A graphically depicts the algorithm’s pipeline. The algorithm is tested in silico by simulating data acquisition using a previously acquired MS dataset and in real-time during an MS experiment.

**Figure 1.**
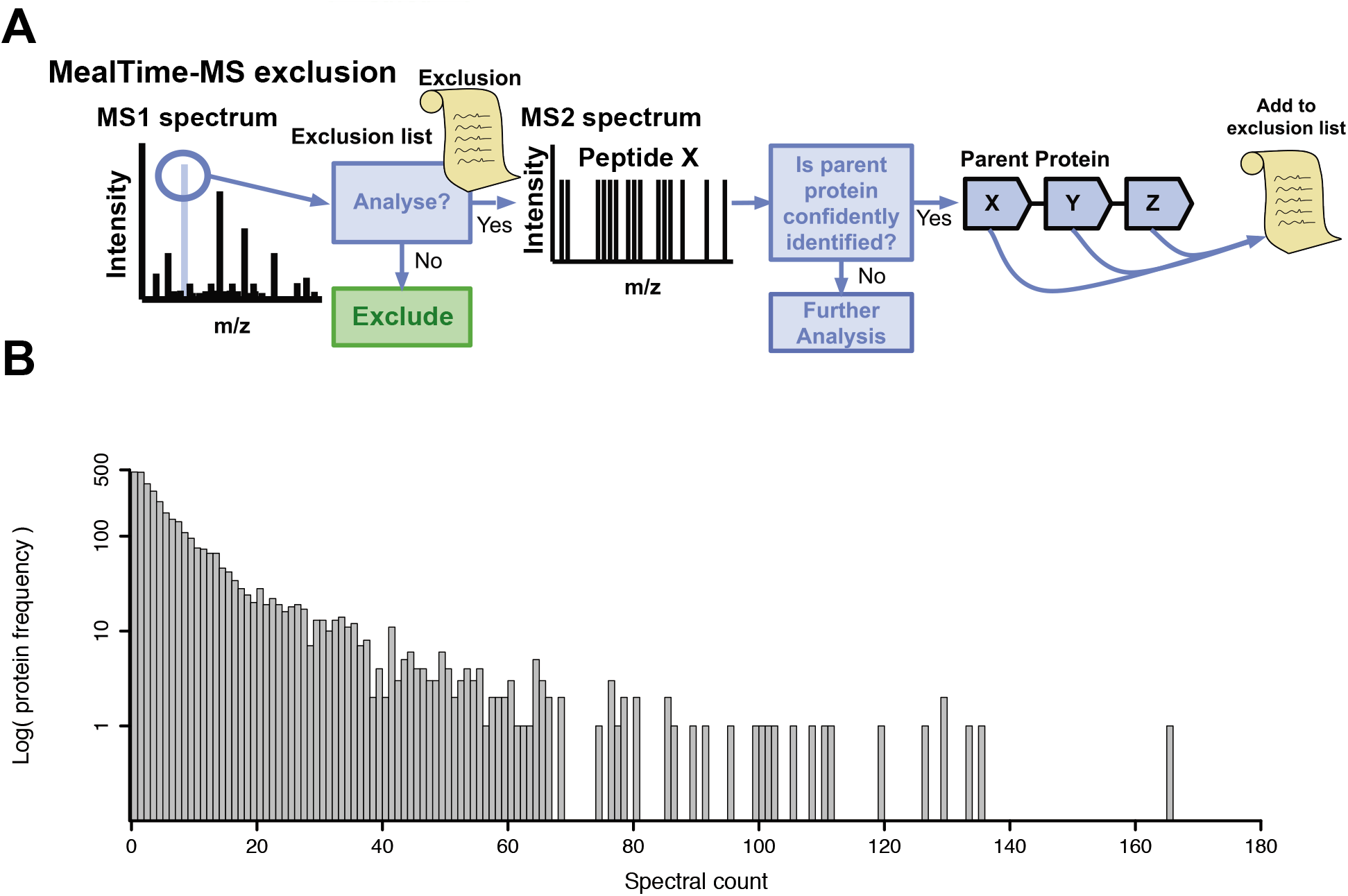
(A) Graphical representation of the MealTime-MS algorithmic workflow. MealTime-MS decides whether an MS2 spectrum should be analyzed or not based on an exclusion list. If analyzed, the confidence of the identification of the parent protein of the peptide identified is assessed using a logistic regression classifier. If the protein is confidently identified, its peptides’ masses are added to the exclusion list. (B) Distribution of the spectral counts of proteins identified (FDR<1%) in the 120-minute HEK293 LC-MS/MS analysis.

### Cell culture and protein extraction

HEK293 cells were grown to 80% confluence in 15 cm dishes and harvested in RIPA buffer (25 mM Tris-HCl (pH 7.6), 150 mM NaCl, 1% NP-40, 1% sodium deoxycholate, 0.5% SDS) after two washes with PBS. Cells were sonicated for 1 min with 20 second pulse using 30% power on Sonic Dismembrator 500 w/ Branson 1020 Sonicator (Fisher Scientific) to increase protein recovery. Proteins were precipitated to remove detergent by adding 5X volume of cold acetone overnight followed by two washes with cold acetone as well.

### Sample digestion

8M Urea in 50mM ABC was used to reconstitute proteins. Reduction and alkylation were done by adding DTT to a final concentration of 10 mM at 56°C for 30 min followed by 20 mM IAA at room temperature. The solution was then diluted 5 times by adding 50 mM ABC. Trypsin was added to achieve a protein-enzyme ratio of 50:1. Digestion was performed at 37°C overnight, with continuous shaking. Digested peptides were then desalted on Sep-Pak column (Waters) and dried down in SpeedVac (ThermoFisher Scientific, San Jose, CA). Dried peptides were reconstituted in 0.5% (v/v) FA, 2% ACN in water.

### LC-MS/MS Analysis

Eksigent 2D+ nanoLC system (Dublin, CA) was hooked up with a Q-Exactive mass spectrometer (Thermo Electron, Waltham, MA), equipped with a nano-electrospray interface operated in positive ion mode. The solvent system consists of buffer A of 0.1% FA in water, and buffer B of 0.1% FA in 80% acetonitrile. Reconstituted peptides were loaded on 200 μm I.D. × 20 mm fused silica pre-column packed in-house with 5 μm ReproSil-Pur C18 beads (100 Å; Dr. Maisch GmbH, Ammerbuch, Germany) at a flow rate of 4 uL/min for 10 min, and analyzed on a 75 μm I.D. × 150 mm fused silica analytical column packed in-house with 1.9μm ReproSil-Pur C18 beads (100 Å; Dr. Maisch GmbH, Ammerbuch, Germany) at a flow rate of 200 nL/min for either 120 min or 240 min to create to different datasets of the HEK293 cell lysate. Gradient elution was set from 5 to 35% buffer B (80% ACN). The spray voltage was set to 2.0 kV and the temperature of the heated capillary was 300 °C. The instrument method consisted of one full MS scan from 300 to 1800 m/z followed by the acquisition of data-dependent MS/MS scan of the 12 most intense ions, a dynamic exclusion repeat count of 1 in 30s, and an exclusion duration of 30s. The full mass was scanned in Orbitrap analyzer with R = 70,000 (defined at m/z 400) for MS1 and 17,500 for MS2. To improve the mass accuracy, all measurements in the Orbitrap mass analyzer were performed with a real-time internal calibration by the lock mass of background ion 445.120025. The charge state rejection function was enabled, and charge states with unknown and single charge state were excluded for subsequent MS/MS analysis. All data were recorded with Xcalibur software (ThermoFisher Scientific, San Jose, CA). Raw outputs from previously completed mass spectrometry experiment were converted to mzML file format using MSConvert from the Trans-Proteomics Pipeline^25^.

### Protein identification for supervised learning algorithm training and benchmarking purposes

In total two 120-minute and one 240-minute LC-MS/MS experiments were performed on HEK293 cell lysates. For supervised learning algorithm training and benchmarking purposes, peptides were identified from the two 120-minute (used for benchmarking) and the 240-minute (used for training) LC-MS/MS mzML files using the Comet protein sequence database search algorithm^26^. Comet search was performed against the UniProt Swiss-Prot human protein sequence database (downloaded on 2017-01-11)^27^. Database search was performed with a precursor ion mass tolerance of 20 ppm, considering only fully tryptic peptides and with a maximum of 1 miscleavage allowed. Carbamidomethylation of cysteine was considered as a fixed modification and methionine oxidation as a variable modification. The resulting files were processed using PeptideProphet and ProteinProphet to assess the confidence of peptide and protein identifications, respectively^28,29^. A protein was deemed confidently identified at a False Discovery Rate (FDR) < 1%.

### Real-time peptide identification and protein identification confidence assessment

MealTime-MS uses a supervised learning approach to determine whether a protein is confidently identified. During an MS/MS experiment, when the mass spectrometer acquires an MS2 spectrum, a Comet database search is performed to identify its matching peptide sequence and a logistic regression classifier estimates the probability that the corresponding protein is confidently identified at this point in the experiment. The Comet search is executed using with single spectrum search module using the same search parameters and database described above to identify the peptide matching the MS2 spectrum with the highest cross-correlation value (XCorr)^26,30^. Let *p* be the parent protein of this highest scoring peptide. The probability that *p* is confidently identified is then assessed by the classifier after each Comet search using the following features, derived from Comet results: 1) the cardinality of the set *S* of MS2 spectra matched so far in the experiment to peptides belonging to *p* (spectral count), 2) the highest XCorr of the peptide-spectrum matches in *S*, 3) the mean and median of the XCorr values of the peptide-spectrum matches in *S*, 4) the highest Delta Cn^30^ value of the peptide-spectrum matches in *S*, and 5) the mean and median of the Delta Cn values of the peptide-spectrum matches in *S*. Using these features, the classifier computes the probability that *p* is reliably identified. A probability threshold *P* determines whether a protein is confidently identified (≥ *P*) or not (< *P*). The logistic regression classifier is trained on the above protein features obtained from the completed 240-minute LC-MS/MS analysis of the HEK293 cell lysate from which positive examples of confident protein identifications and negative examples of proteins that have not been identified with confidence were extracted. PeptideProphet and ProteinProphet were used to assess the confidence of a protein identification obtained using Comet^28,29^. Proteins identified with a FDR < 1% according to ProteinProphet were used as positive examples of high confidence protein identifications. All other proteins, including decoy proteins were used as negative examples of protein identifications. In total, 3796 positive and 9515 negative examples were used for training with the iteratively reweighted least squares algorithm^31^. The positive training set for a confident protein identification was chosen using a 1% FDR threshold since it is a standard in the field for a high-confidence protein identifications. The resulting real-time confident identifications of proteins are then used to build and maintain a dynamic exclusion list of peptide masses.

### Peptide Retention Time Prediction and Correction

Since multiple distinct peptides in a complex sample, such as a cell lysate, can have similar or equal masses and m/z values, it is important to add a given mass to the dynamic exclusion list only during the period of time at which the peptide is expected to elute during the MS experiment. We call this period a mass exclusion time window. To build such a window, the peak retention time of each peptide was predicted using RTCalc from the Trans-Proteomics Pipeline and by adding a constant *w* in minutes before and after the predicted retention time^25^. Since peak peptide retention time can vary due to experimental conditions, if a peptide is matched to an MS2 spectrum with a XCorr score > 2.5 (strong match^32^) during the MS/MS analysis, its mass exclusion time window is adjusted such that the time this peptide was observed by the mass spectrometer corresponds to the beginning of the time window. Additionally, the time difference between a peptide’s observed and predicted retention time is also used to calibrate the predicted retention times of all peptides in real-time. For each peptide with a XCorr score > 2.5, MealTime-MS calculates the difference between the peptide’s predicted peak retention time and the time at which this peptide is observed. This time difference is then used to calculate a time offset using the *R* most recent differences. The offset is calculated using a convergent geometric series that weighs the differences in retention times in favor of the more recent values:

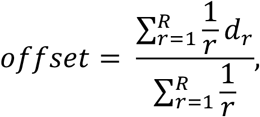

where *d_r_* is the time difference of the r^th^ peptide from *R*. The offset is then added to the RTCalc predicted retention time to more accurately reflect when a peptide might be observed in the MS experiment.

### Real-time dynamic exclusion list construction and maintenance

MealTime-MS leverages the dynamic exclusion list used in Top N DDA MS/MS. The mass spectrometer uses such a dynamic exclusion list, consisting of peptide masses it will not select for fragmentation and MS2 spectrum acquisition. Ions with a m/z value within a given mass tolerance *m* measured in parts per million (ppm) of an excluded mass on the exclusion list (after single charge transformation) will not be selected for fragmentation. Top N DDA will therefore acquire MS2 spectra for the N most intense ions that are not excluded. In MealTime-MS, the dynamic exclusion list is constructed as the algorithm processes MS2 spectra and assesses the confidence of protein identifications. Peptides that are matched with an MS2 spectrum with a XCorr score > 2.5 and peptides from proteins identified with high confidence (logistic regression probability ≥ *P*) will see their masses added to the dynamic exclusion list in real-time by our software. The latter peptides are excluded based on a peptide sequence database. This database is generated in silico prior to the MS run using the UniProt Swiss-Prot sequence database (downloaded on 2017-01-11)^27^ and digested in silico using the chainsaw tool from the ProteoWizard suite with trypsin as the digestion enzyme parameter^33^. Peptides generated are fully tryptic, have at most one miscleavage, and have a minimum and a maximum length of 6 and 200 amino acids, respectively. Their masses are calculated using their amino acid constitution. Hence, once a protein is confidently identified by MealTime-MS, the masses of all peptides associated to it in the peptide sequence database are added to the exclusion list for the period defined by its mass exclusion time window (described in previous section).

During certain time periods of the MS experiment, it is possible that MealTime-MS will not be able to process MS2 spectra as fast as they are acquired. To prevent the loss of MS2 spectra that are received by MealTime-MS before it was done processing preceding MS2 spectra, acquired mass spectra are sent into a first in first out queue, where they wait until being processed. To prevent accumulating a large queue where MS2 spectra would be analyzed long after they were acquired, the queue size is capped to 25. Once it reaches this limit, MS2 spectra are ignored by MealTime-MS until the mass spectra in the queue are processed such that less than 10 remain, after which the queue resumes taking MS2 spectra in.

### Simulating mass spectrometry analysis in silico

While we tested MealTime-MS in real-time during one of the two 120-minute LC-MS/MS experiments, the majority of our benchmarking, parameter testing and performance evaluations were performed using in silico simulations of mass spectrometry data acquisition using the other 120-minute LC-MS/MS analysis, which was previously completed. In these simulations, the algorithm processes mass spectra from the mzML file in the order they were previously acquired by the instrument. It also simulates the time delays between each mass spectrum acquisition based on their recorded time during the original MS experiment. Before processing a mass spectrum, MealTime-MS makes a real-time decision based on the dynamic exclusion list whether to perform a database search on that mass spectrum and assess the confidence of the putative protein identification or to ignore it to simulate its exclusion. Based on the machine learning protein confidence assessment our tool updates the dynamic exclusion list. To assess the number of proteins identified, our software outputs a database search result file of the MS2 spectra that were not excluded from MS/MS analysis (pep.xml file), such that PeptideProphet and ProteinProphet can be executed. At the same time, the software package keeps track of the MS2 spectra that were excluded from the analysis and not used for protein identification. These are considered saved MS2 spectra that could be redistributed to acquire MS2 spectra from proteins that remain uncharacterized.

While the algorithm was trained on a 240-minute LC-MS/MS experiment, an assumption is made that a confident protein identification from a 240-minute experiment will share similar properties to a confident protein identification from a 120-minute experiment. Training our algorithm on an experiment using a different LC gradient length also showcases the ability of our approach to be generalizable to different datasets. Nevertheless, the datasets used in this study remain similar, since they analyze the same cell type and are processed with the same MS instrument.

### Heuristic approach for protein identification confidence assessment

MealTime-MS performances were benchmarked against a heuristic-based approach inspired by a method proposed by McQueen et al.^21^ The heuristic exclusion strategy matches the implementation of our machine learning-guided dynamic exclusion in all aspects of the algorithm with the exception of its protein identification confidence assessment method. With the heuristic approach, a protein *p* is deemed confidently identified when a count of *s* MS2 spectra associated with *p*, are matched with peptides of *p* with a XCorr score > *x*.

### Implementation and availability

MealTime-MS is implemented as an open-source C# program. All code and documentation have been deposited at the following GitHub repository: https://github.com/LavalleeAdamLab/MealTimeMS.git

MealTime-MS uses the Thermo Fisher Instrument Application Programming Interface (IAPI) to collect MS data in realtime and in our simulations. The IAPI is available here: https://github.com/thermofisherlsms/iapi. Only the open version of the IAPI was used in this study. The exclusion list built by MealTime-MS was therefore not transferred to the mass spectrometer during real-time analysis. MS2 spectra of ions that matched the exclusion list were therefore acquired by the instrument, but ignored by MealTime-MS to simulate exclusion. The MS-based proteomics data have been deposited to the ProteomeXchange Consortium^34^ via the PRIDE^35^ partner repository with the dataset identifier PXD017673.

## RESULTS

MealTime-MS is a machine learning-based approach assessing protein identification confidence in real-time during MS experiments. Based on the peptides associated with confidently identified proteins, it updates a dynamic exclusion list to prevent the redundant acquisition of mass spectra related to these proteins. Figure 1A graphically depicts MealTime-MS’ pipeline. The algorithm is tested using in silico simulations of MS data acquisition from a previously obtained proteomics dataset and in real-time during an actual MS experiment. Both datasets were analyzed using a Top 12 DDA of HEK 293 cell lysates.

### A large number of mass spectra acquired in a typical proteomics experiment do not favor protein identification

To evaluate the potential gain that could be achieved by preventing the acquisition of MS2 spectra for peptides related to already confidently identified proteins, we investigated the spectral count distribution of confidently identified proteins of a previously acquired MS dataset using a Top 12 DDA and standard dynamic exclusion method (see Methods). Figure 1B shows the distribution of the spectral counts for proteins identified (FDR < 1%) in a 120-minute LC-MS/MS analysis of a HEK 293 cell lysate. Assuming a very conservative spectral count threshold of 5 MS2 spectra necessary to identify a protein, 1,541 proteins would be detected in this dataset (Figure 1B). From those proteins, 61.1% of the MS2 spectra would be in excess of what is necessary for protein identification if 5 mass spectra were required. This illustrates that the majority of MS2 spectra acquired in this experiment could have been used more efficiently to acquire data from poorly detected or uncharacterized proteins.

### Retention time calibration improves retention time prediction

Since peptides from proteins deemed confidently identified by MealTime-MS will see their masses added to the dynamic exclusion list for a given mass exclusion time window, RTCalc was used to determine the expected peak retention time of peptides added to the exclusion list. However, accurate peptide retention time prediction in LC is challenging and predictions for our 120-minute LC-MS/MS experiment were initially quite poor. Supplementary Figure S1A shows the retention time prediction errors for all observed peptides. The error was calculated as the difference between a peptide’s predicted retention time and the experimental time at which this peptide was detected by the mass spectrometer. Only 0.3% of the peptide-spectrum matches were detected within 1 minutes of their peptide predicted retention time, with some of the retention time errors reaching more than 100 minutes (Supp. Fig. S1A). Our software package therefore corrects mass exclusion time windows based on the observed retention time. An offset is also computed to calibrate retention time predictions using the *R* most recent differences between a peptide’s predicted and observed retention times of peptide-spectrum matches with XCorr score > 2.5 (see Methods). The performance of this calibration was assessed by comparing the fraction of peptides whose actual retention times fell within their predicted retention time windows (predicted retention time ± *w* (in minutes)). Incorporating a greater number of *R* error values in the offset calculation slightly increased the number of peptides with retention times falling within *w* of the predicted retention time (Supp. Table S1), with the exception of *R* = 5000 that saw a decrease. As expected, when increasing *w*, a greater percentage of peptides fell within their predicted retention time window. *R* = 100 real-time MS analyses since it yielded a percentage of observed peptides falling within one minute of their predicted retention times that was similar to other *R* values and it allows the offset to be computed more rapidly. Calibration with *R* = 100 increased the percentage of peptides within 1 minute of their predicted retention times from 0.3% to 29.6% (~100-fold) (Supplementary Figures S1A and S1B).

For peptide retention time calibration to be achieved on-the-fly, the offset values must be calculated only from MS2 spectra acquired by the mass spectrometer. Since MealTime-MS excludes from MS/MS analysis a number of these mass spectra, we assessed the performance of the peptide retention time calibration using only the MS2 spectra that were not excluded using our algorithm. The 120-minute LC-MS/MS analysis was simulated with machine learning-guided exclusion with on-the-fly retention time calibration. Masses of peptides from proteins deemed confidently identified (identification probability threshold P = 0.5) were excluded, with a precursor mass tolerance *m* = 5 ppm, for *w* =1 minute around their offset-corrected predicted peptide retention time. With this approach, 0.45% of peptide-spectrum matches fell within 1 minute of their predicted retention time before retention time calibration (Supp. Fig. S1C), compared to 14.8% with calibration (Supp. Fig. S1D).

### MealTime-MS yields a high protein identification sensitivity despite excluding large amounts of MS2 spectra from the analysis

To assess the performance of our algorithm, we simulated MS data acquisition on a previously completed 120-minute LC-MS/MS analysis of a HEK293 cell lysate, processing the mass spectra in the order they were originally acquired. This experimental simulation was repeated using a variety of algorithm parameters: *w* = 0.75, 1, and 2 minutes, *m* = 5 and 10 ppm, and *P* = 0.3, 0.5, 0.7, and 0.9. Our logistic regression classifier was benchmarked against our heuristic exclusion approach, inspired by the work of McQueen et al.^21^, for assessing the identification of a protein *p* based on identifying *s* MS2 spectra matched to peptides of *p* with a XCorr > *x*, where *s* was set to 1 or 2 and *x* to 1.5, 2, or 2.5 (see Methods). The simulations using the heuristic approach were performed with the same values of *m* and *w* described above for MealTime-MS for the exclusion list. In addition to this benchmarking, we evaluated how these two algorithmic approaches fared against a random exclusion strategy. A subset of the MealTime-MS and heuristic-guided simulations were matched by a set of ten simulations, where the same number of MS2 spectra the two algorithms excluded were excluded at random. Figure 2 and Supp. Table S2 show the results from the simulations of the 120-minute LC-MS/MS experiment with the above algorithms and parameters. The combination of the following parameters: *m* = 5 ppm, *w* = 1 minute and *P* = 0.5, yielded one of the best simulation results for MealTime-MS, which identified 92.1% of the proteins detected in the original experiment (protein identification sensitivity), while using only 66.2% of its MS2 spectra. Both MealTime-MS and the heuristic exclusion approaches obtained a greater protein identification sensitivity than the random exclusion for the same number of exclude MS2 spectra, indicating that the exclusions made by these algorithms were less deleterious than those performed by the random strategy. However, when roughly using the same number of MS2 spectra for protein identifications (i.e. not excluded MS2 spectra), MealTime-MS protein identification sensitivity is on par with the one from the heuristic approach. Since both approaches do not exclude the exact same set of MS2 spectra from MS/MS analysis, we developed a combined approach, where an MS2 spectrum would be excluded if one of the two methods would do so. Interestingly, this combined approach yields only marginal improvements in terms of protein identification sensitivity over both the MealTime-MS method and the heuristic approach for the same number of MS2 spectra used. While this result is somewhat surprising, we hypothesized that it stemmed from the fact that since retention time predictions are relatively unreliable, the time at which a peptide is excluded is often wrong, yielding little to no benefits of performing additional exclusions. To validate this claim, we performed another set of simulations, where we replaced RTCalc retention time predictions for each peptide by its actual measured retention time in the experiment with a random perturbation of ± 15 seconds. Peptides from proteins that were confidently identified were therefore added to the exclusion list using a mass exclusion time window much closer to the actual time they eluted in the experiments. This modification drastically improved the protein identification sensitivity for all approaches. Although, this time, MealTime-MS showed a higher protein identification sensitivity than the heuristic approach for the same number of MS2 spectra used. The combined approach also showed clear improvements over the heuristic approach with only a marginal gain over MealTime-MS alone, showing that the heuristic approach only identifies a very small set of proteins that MealTime-MS does not using a similar number of MS2 spectra. MealTime-MS reached a protein identification sensitivity of 97.9% using only 59.2% of the MS2 spectra with *m* = 5 ppm, *w* = 1 minute and *P* = 0.5, a significant improvement over its previous results using RTCalc retention time predictions (Figure 2 and Supp. Table S2). Of note, both MealTime-MS and the heuristic approach reached an almost perfect protein identification sensitivity (99.5% and 99.2%), while using only 72.4% and 72.1% of the MS2 spectra from the original experiment, respectively (Figure 2 and Supp. Table S2). In an actual experiment, unused MS resources by MealTime-MS could be used to acquire MS2 spectra corresponding to lower abundance proteins and therefore help to characterize more proteins in a biological sample.

**Figure 2.**
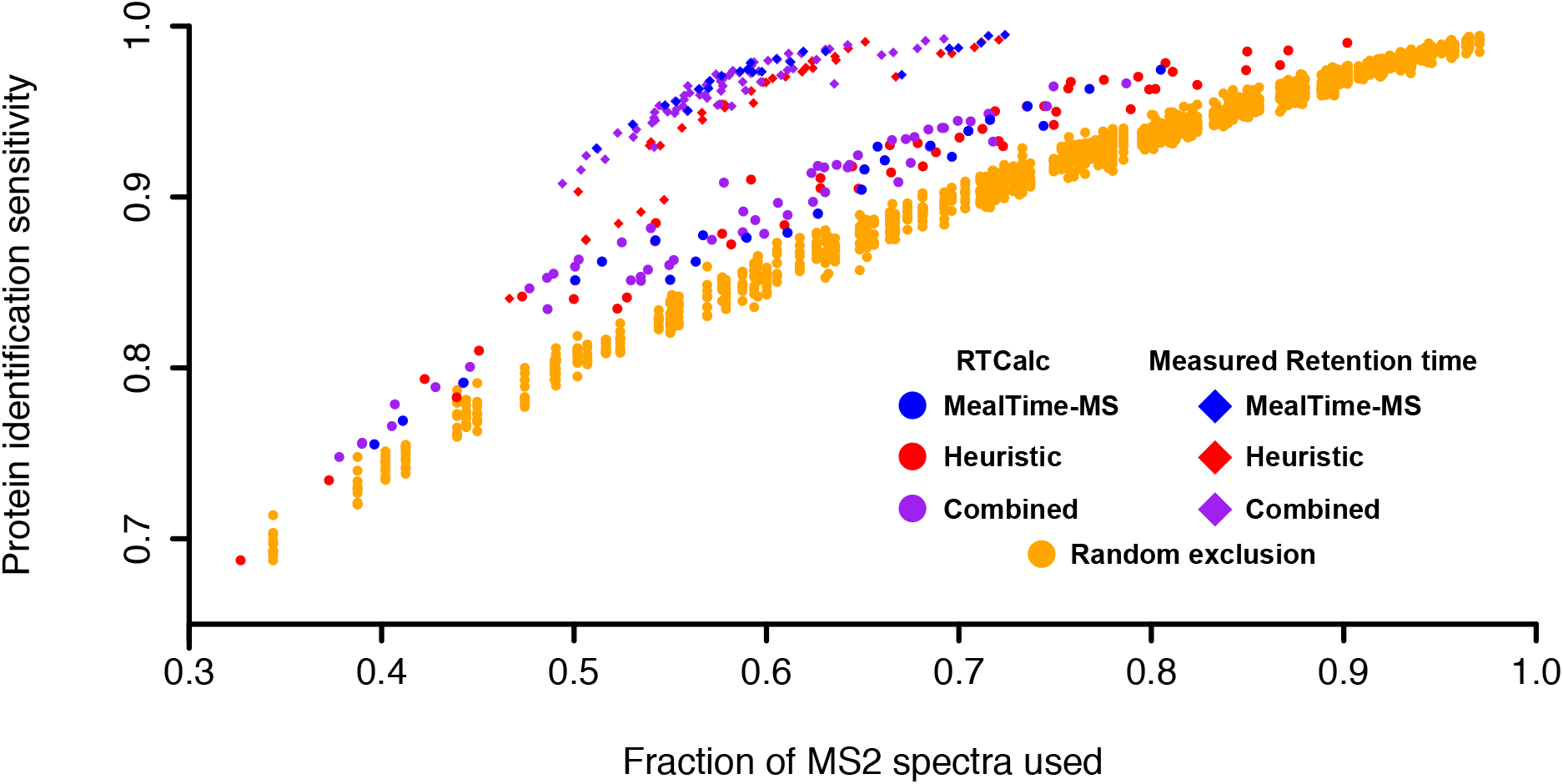
Fraction of MS2 spectra used and protein identification sensitivity for simulations using the MealTime-MS (blue), heuristic (red), combined (purple) and random (orange) real-time exclusion algorithms. MealTime-MS, heuristic, and combined algorithms are used with retention time predictions from RTCalc (circles) and from previously measured retention time with mild perturbation (diamonds). Performances of all algorithms are assessed using the fraction of MS2 spectra used (i.e. not excluded) for protein identification and the fraction of proteins identified (FDR< 1%) using only non-excluded mass spectra over the total number of proteins previously identified (FDR< 1%) using all available spectra.

### Most MealTime-MS exclusions from fragmentation are correct

Next, we assessed the correctness of the machine learning algorithm’s peptide exclusion with P = 0.5. A correct exclusion is defined as an MS2 spectrum that is excluded because its corresponding peptide has indeed been previously added to the exclusion list. An incorrect exclusion on the other hand corresponds to an MS2 spectrum for which the corresponding peptide was not on the exclusion list, but its precursor ion mass was within the ppm mass tolerance of a mass from another excluded peptide, while sharing similar retention time predictions. Figure 3 depicts the number of correct and incorrect exclusions of MS2 spectra. As expected, increasing the ppm mass tolerance for the masses on the exclusion list allows the exclusion from fragmentation of more ions. A larger mass tolerance is also affiliated to an increase in incorrect exclusions with 10 ppm showing the highest incorrect exclusion rate for similar total number of exclusions (Figure 3). On the other hand, increasing the mass exclusion time window size of the masses on the exclusion list resulted in more spectra being excluded overall, with both increased correct and incorrect exclusions (Figure 3). Nevertheless, Figure 3 demonstrates that in the vast majority of simulations, MealTime-MS performed more correct than incorrect exclusions, despite using poor retention time predictions. Correct and incorrect exclusions varied based on the stringency of the algorithm parameters, with a smaller ppm mass tolerance resulting in fewer incorrectly excluded spectra than a large ppm mass tolerance.

**Figure 3.**
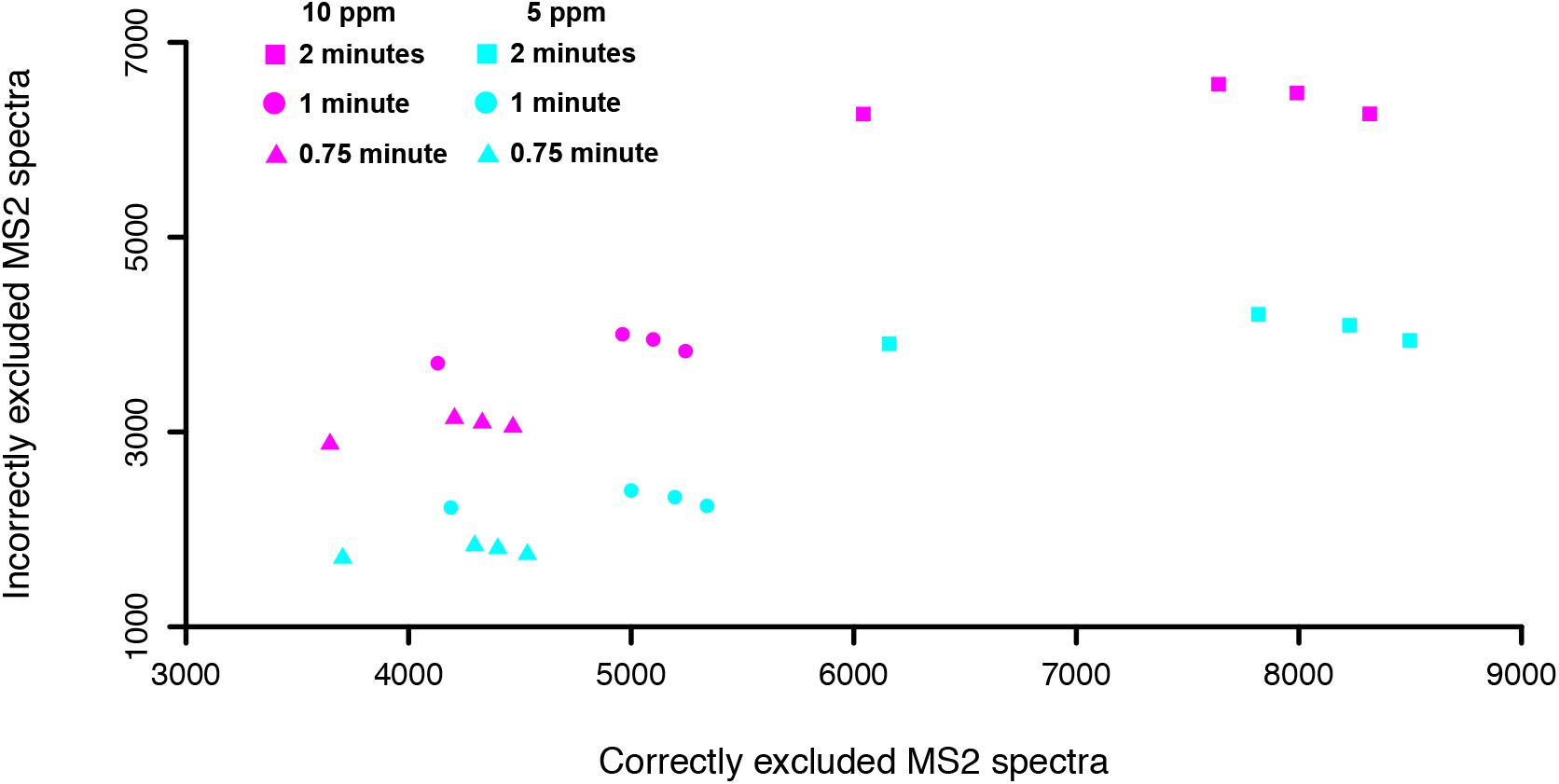
Number of correctly and incorrectly excluded MS2 spectra using MealTime-MS with different algorithm parameters. Mass tolerance *m:* 5 ppm (cyan) and 10 ppm (magenta). Mass exclusion time window *w*: 0.75 minute (triangle), 1 minute (circle), and 2 (square) minutes and protein identification probability threshold *P*: 0.3, 0.5, 0.7, and 0.9.

### MealTime-MS excludes from fragmentation peptides from a wide array of proteins

To ensure that MealTime-MS was not preventing MS2 spectra acquisition for peptides of only a few abundant proteins, we investigated the spectral counts of the proteins for which peptides would be denied fragmentation by MealTime-MS. The opposite behavior would indicate that a similar performance to MealTime-MS could be achieved merely by arbitrarily excluding a handful of highly abundant proteins. Figure 4 shows the number of MS2 spectra that were excluded by MealTime-MS and the randomized exclusion for each protein in the 120-minute experiment with *m* = 5 ppm, *w* = 1 minute and *P* = 0.5. While proteins with high spectral counts generally saw more of their MS2 spectra excluded by our approach, 42.7% of all putative proteins detected by MS have seen at least one of their MS2 spectra excluded. This illustrates that our exclusion strategy does not derive a vast majority of its saved resources from a handful of highly abundant proteins but appears to exclude peptides from a wide array of proteins detected in the experiment.

**Figure 4.**
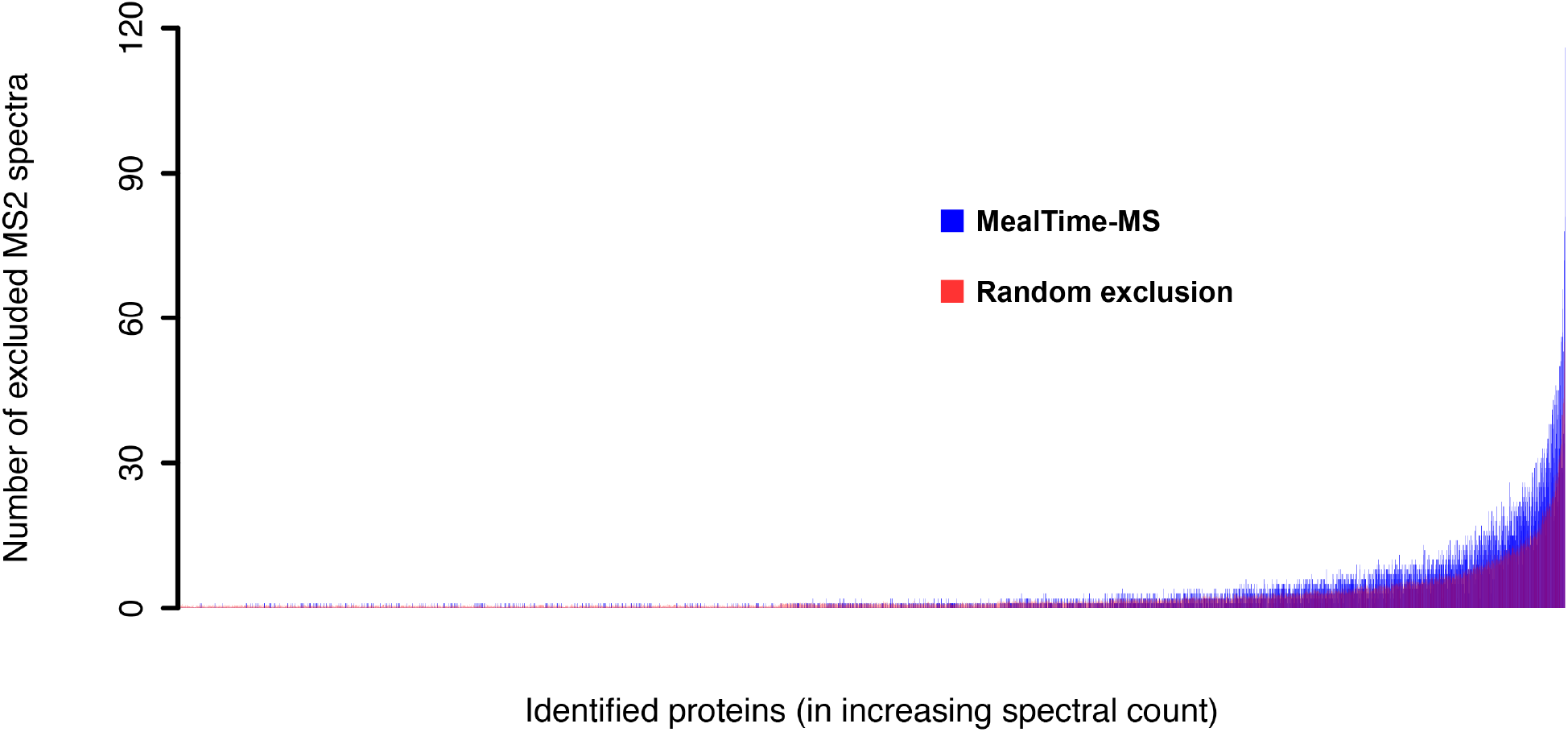
Number of excluded MS2 spectra from detected proteins using MealTime-MS and random exclusion approaches for the 120-minute LC-MS/MS dataset. Peptides from proteins achieving a protein identification probability above 0.5 were excluded for 1 minute around their predicted peak retention times. MS2 spectra with a precursor mass within 5 ppm of a mass on the exclusion list were excluded. Detected proteins on the x-axis are sorted in ascending order of abundance based on their spectral counts in the original experiment. Numbers of excluded MS2 spectra for the random exclusion correspond to the average of 10 randomized runs of the exclusion strategy.

### MealTime-MS is fast enough to perform real time mass spectrometry analysis

Finally, for MealTime-MS to be used during a MS experiment, it must be able to assess protein identification confidence and update the instrument’s dynamic exclusion list within a reasonable timespan, ideally as fast as spectrum acquisition speed. Figure 5A shows our algorithm overall running time for simulations under the different values of *m*, *w* and *P* listed above. As expected, simulations with more lenient parameters resulted in a larger number of peptides added to the exclusion list, thereby increasing the running time of the algorithm. Depending on the number of peptides added to the exclusion list, our algorithm takes from 1,122 to 3,141 seconds of total computational time to assess protein identification confidence, decide whether a MS2 spectra should be excluded or processed, and update the dynamic exclusion list. All instances of the simulations took well below the 7,200 second duration (120 minutes) of the MS analysis. Nevertheless, mass spectra are not acquired uniformly throughout those 120 minutes. Certain periods of the experiments generate large amounts of mass spectra and put a heavier load on MealTime-MS. Furthermore, our simulations did not consider the network latency involved with the mass spectrometer and the supporting computer running MealTime-MS communicating with each other. We therefore also ran MealTime-MS during an actual MS experiment analyzing a HEK293 cell lysate for 120 minutes using *m* = 5 ppm, *w* = 1 minute, and *P* = 0.5. Figure 5B and 5C show that during this real-time analysis 24.9% of MS2 spectra were processed (analyzed or excluded), within 10 milliseconds or less (including the time spent in the queue), which is below most mass spectrometer’s rate of acquisition. In addition, more than 75% of the MS2 spectra were processed in less than 0.1 seconds. Our queue approach for MS2 spectra processing led to only 1.46% spectra to be completely ignored by MealTime-MS in order to prevent the software from lagging behind MS acquisition (Figure 5B). Figure 5B highlights that shortly after ignoring MS2 spectra, MealTime-MS processing time rapidly reduces. These results show that the information provided by MS2 spectra can be rapidly used by MealTime-MS to enrich its decision process for future exclusions during an MS/MS experiment.

**Figure 5.**
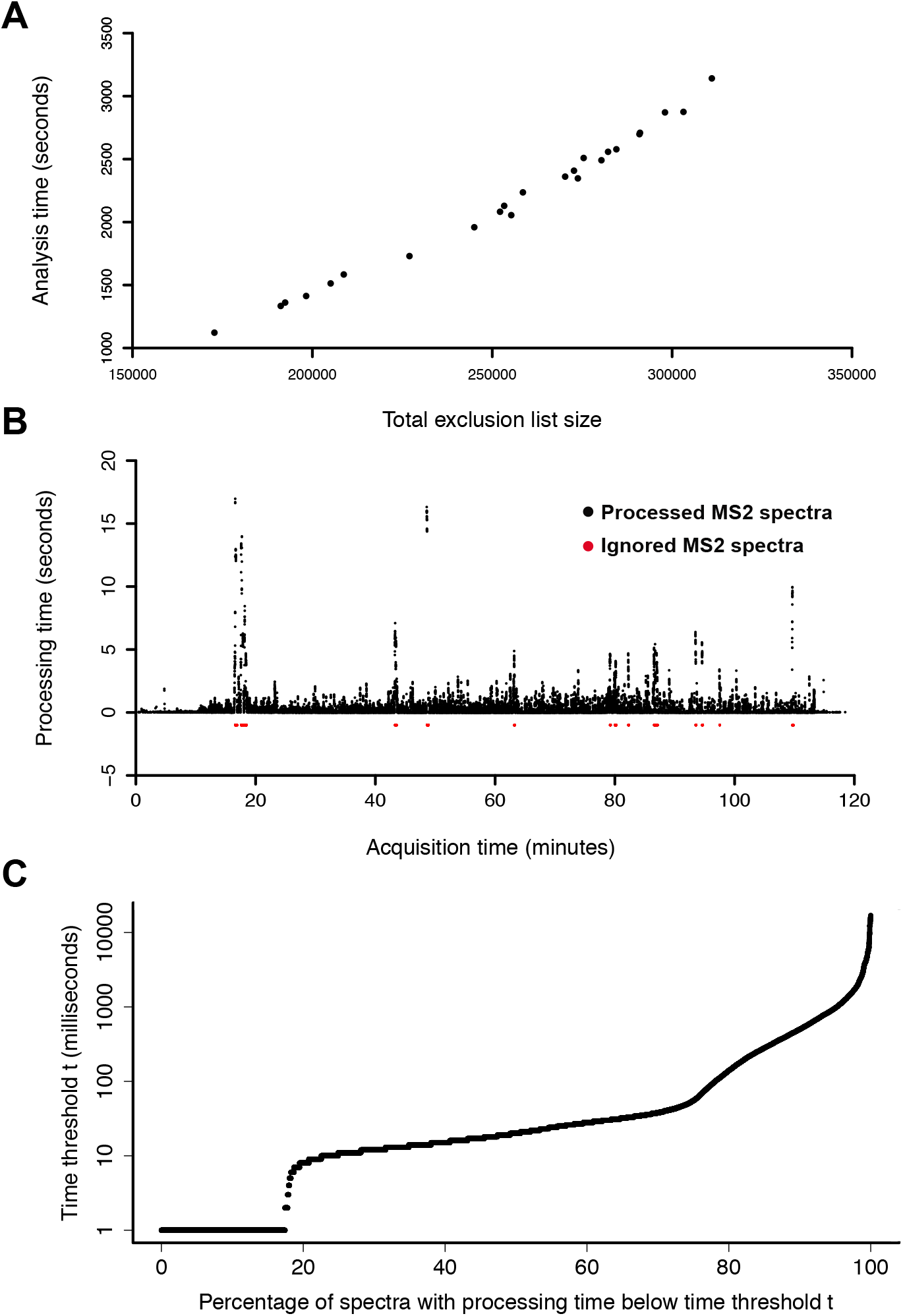
Execution time analysis of MealTime-MS. (A) Analysis time of MealTime-MS for simulations performed on the previously acquired 120-minute LC-MS/MS analysis as a function of the total number of peptides added to the exclusion list during the simulation. Simulations were run for ppm mass tolerance *m*: 5 and 10 ppm, mass exclusion time window *w*: 0.75, 1, and 2 minutes, and protein identification probability threshold *P*: 0.3, 0.5, 0.7, and 0.9. (B) Time taken by MealTime-MS to process MS2 spectra in real-time on a desktop computer connected to the mass spectrometer while it was acquiring data. The algorithm was executed m = 5 ppm, w = 1 minute, and P = 0.5. Red points indicate MS2 spectra that were ignored by MealTime-MS when the analysis queue was full, while black points represent processed MS2 spectra. (C) Percentage of MS2 spectra that were processed within a given time threshold.

While MealTime-MS is a computationally demanding piece of software, we tested the package to ensure that it could be executed on a regular desktop computer, such as those supporting most MS instruments during data acquisition. Our real-time analysis from Figure 5B and 5C was performed on an Intel Core i3-2120 (3.30 GHz) computer with 4 GB of RAM running a 32-bit Windows operating system, which was connected to the mass spectrometer. This demonstrates that the software package can process most mass spectra rapidly without any exceptional computing hardware. Simulations were performed on an Intel Core i7 6700 (3.4 GHz) desktop with 16 GB of RAM running a 64-bit Windows operating system and an Intel Core i5-6600K (3.5GHz) desktop with 8 GB of RAM running a 64-bit Windows operating system.

## DISCUSSION

### Retention time prediction plays a key role in the success of real-time MS analysis

Our simulations show that the more accurate retention time predictions are, the greater MealTime-MS performances will be. Variations in samples and LC conditions can drastically affect our ability to predict peptide retention times. As an alternative to provide better retention time predictions, samples could be run in duplicates, where one replicate could serve as a retention time reference to provide better predictions. Nevertheless, such an approach requires more samples and analytical time and defies the purpose of MealTime-MS, which attempts to identify proteins using less mass spectra. Furthermore, results from other retention time prediction tools for LC, such as SSRCalc^36^ or DeepRT^37^ could be used to potentially generate a better exclusion list. Results from these tools could be integrated along with RTCalc to generate a consensus retention time prediction that may be more representative of actual peptide retention times.

### Machine learning classifier training and parameter adjustments

We showed that MealTime-MS can be trained on an experiment using a given LC gradient and be used on an experiment using a gradient of a different duration. While a single dataset seems sufficient to train our classifier, it is possible that it would benefit from being trained on a set of experiments in order to be generalizable to different instruments and samples. Nevertheless, an advantage of this machine learning approach is that training requires very few parameter selections, unlike the heuristic approach, where one has to balance out hard thresholds on the number of mass spectra required for a confident protein identification and their minimal matching score according to a database search algorithm. Choosing these parameters can be difficult, since accepting protein identifications with a single peptide-spectrum match can be sometimes too lenient depending on the matching score, while requiring two peptide-spectrum matches can be too stringent in other cases. With MealTime-MS, proteins with both one or two peptide-spectrum matches can be both a confident identification or low reliability one. The only parameter the user has to select is the threshold *P* for the output probability of the classifier, which controls the confidence in a protein identification.

### Repurposing excluded MS2 spectra

In its current state, MealTime-MS is only capable of determining, which ions should be excluded from fragmentation. At this time, most mass spectrometry vendors’ APIs do not openly provide a full control over their instruments. The IAPI used in this work is open-source, freely available and allows real-time access to mass spectra acquired by the mass spectrometer. However, functionalities allowing third party software to send commands to the mass spectrometer, such as updating an exclusion list or acquiring a mass spectrum from a given ion instead of another are locked unless a license key is provided by the vendor with which an agreement must be made. With such functionalities, the MS2 spectra excluded by MealTime-MS could be repurposed to characterize peptides of lower abundance and therefore provide a more in depth coverage of the proteome of a sample. While we applaud the increasing availability of such IAPIs, only when these tools will be fully open and allowing a large community of computer scientists to develop new approaches will the field of real-time MS analysis truly impact proteomics research.

### Impacts on protein quantification

Since MealTime-MS can alter the number of MS/MS spectra that are acquired for a given protein based on real-time protein identifications, label-free quantification of peptides or proteins using spectral counts^1^ becomes unreliable with this strategy. The goal of MealTime-MS is not to quantify proteins. Nevertheless, MS1 spectrum-based label-free quantification, such as Extracted Ion Chromatogram (XIC) could still be performed and would not be impacted by our procedures. Similarly, stable isotope labeling approaches would not be impacted, with the exception that potentially less MS/MS spectra would be acquired for high abundance peptides labeled using TMT^38^ or iTRAQ^39^.

### Incorrect exclusions do not need to be avoided at all costs

While we showed that MealTime-MS does make a significant number of incorrect exclusions, the goal of our approach should not be to eliminate all of them at the detriment of excluding less MS2 spectra. Indeed, if an MS2 spectrum is reattributed to a different ion it was originally intended for, it will still contribute to a protein identification and is therefore not wasted. However, such an MS2 spectrum would be used on an ion that is most likely of lower intensity and could therefore be of lesser quality and more difficult to identify than the one that would have been generated by the original precursor ion.

## CONCLUSIONS

In conclusion, MealTime-MS will contribute to paving the way to the development of new machine learning-based methods for real-time MS analysis. This work also presents a framework to test real-time MS analysis methods using simulations. Furthermore, in the future, it will contribute to provide a more comprehensive proteomic characterization of complex samples with proteins of high abundance, such as those originating from blood or microbiota, all of which while using fewer mass spectrometry resources.

## ASSOCIATED CONTENT

### Supporting Information

The Supporting Information is available free of charge on the ACS Publications website.

Supplementary Figure S1. Retention time prediction error before and after retention time calibration (PDF).

Supplementary Table S1. Impact of retention time calibration on prediction accuracy (XLSX).

Supplementary Table S2. Results from the simulated real-time MS analysis using MealTime-MS, heuristic, combined and random algorithms under varying parameters (XLSX).

## AUTHOR INFORMATION

### Author Contributions

The manuscript was written through contributions of all authors. / All authors have given approval to the final version of the manuscript. /

## ACKNOWLEDGMENT

The authors acknowledge funding from the following sources: Natural Sciences and Engineering Research Council of Canada Discovery grants to M.L.A. and D.F. Funding from the Government of Canada through Genome Canada and the Ontario Genomics Institute (OGI-156), the Natural Sciences and Engineering Research Council of Canada (NSERC, grant no. 210034), and the Ontario Ministry of Economic Development and Innovation (ORF-DIG-14405) to D.F. A.R.P was funded by a stipend from the NSERC CREATE in Technologies for Microbiome Science and Engineering (TECHNOMISE) Program.

