## Supplementary Figure S1 for "MealTime-MS: A Machine Learning-Guided Real-Time Mass Spectrometry Analysis for Protein Identification and Efficient Dynamic Exclusion"

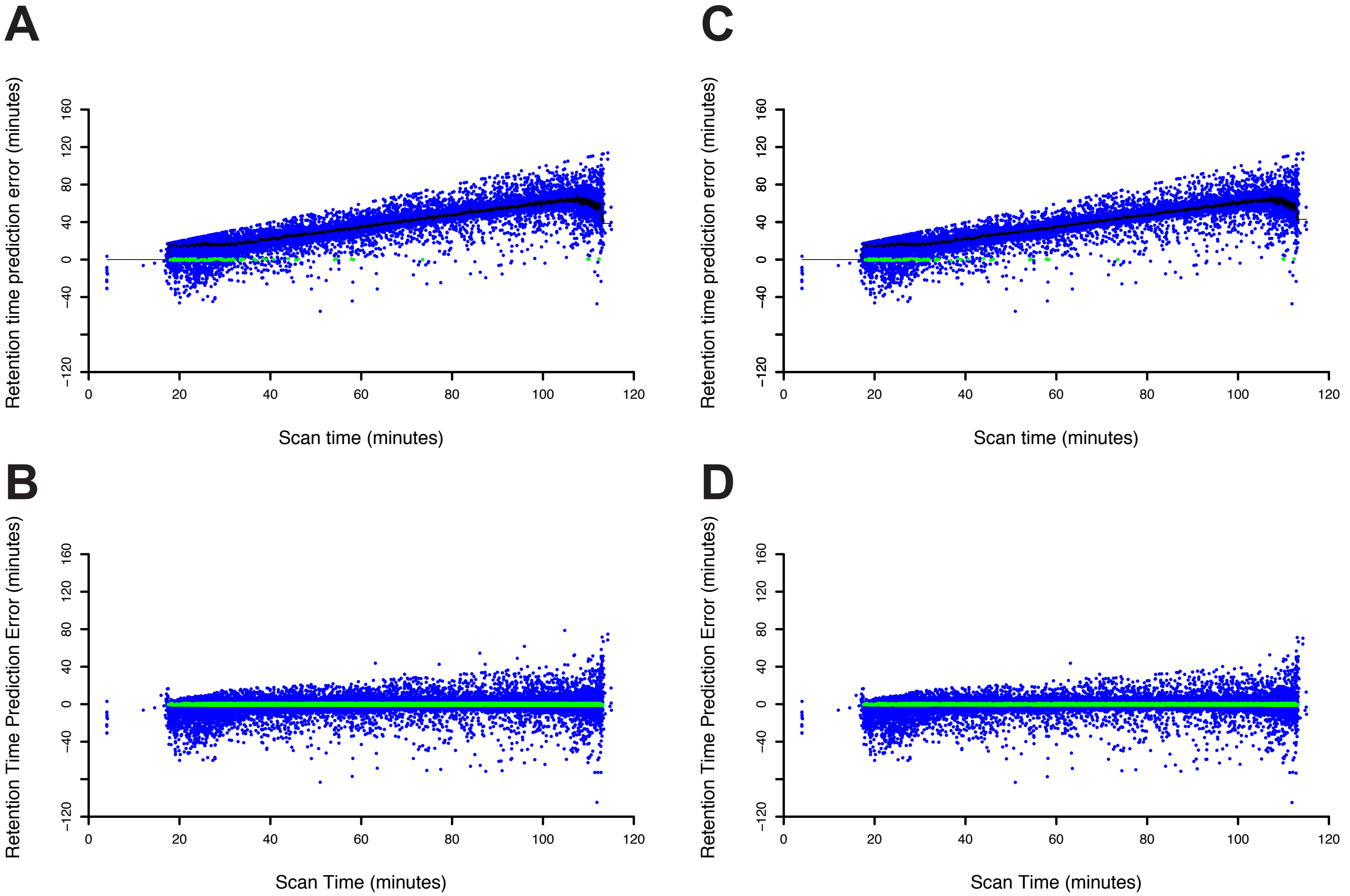


**Supplementary Figure S1. Retention time prediction error before and after retention time calibration.** (A) Error from RTCalc retention time prediction is shown for each peptide-spectrum matches of the 120-minute LC-MS/MS analysis at its experimental time. Peptides detected during the experiment within 1 minute of predicted retention time are shown in green. The offset calculated using the weighted mean of the 100 (R) most recent retention time prediction errors is shown in black at each time step of the experiment. (B) Error with calibrated retention time predictions using the offset. (C) Error from RTCalc retention time prediction is shown for peptide-spectrum matches from the same dataset, but only of peptides that were not excluded by MealTime-MS (parameters: m = 5ppm, w = 1 minute and P = 0.5). (D) Error with calibrated retention time predictions using the offset and only not excluded peptides .
